# Size control in mammalian cells involves modulation of both growth rate and cell cycle duration

**DOI:** 10.1101/152728

**Authors:** Clotilde Cadart, Sylvain Monnier, Jacopo Grilli, Rafaele Attia, Emmanuel Terriac, Buzz Baum, Marco Cosentino-Lagomarsino, Matthieu Piel

**Author notes:** Correspondence should be sent to and (lead contact).

## Abstract

Despite decades of research, it remains unclear how mammalian cell growth varies with cell size and across the cell division cycle to maintain size control. Answers have been limited by the difficulty of directly measuring growth at the single cell level. Here we report direct measurement of single cell volumes over complete cell division cycles. The volume added across the cell cycle was independent of cell birth size, a size homeostasis behavior called “adder”. Single-cell growth curves revealed that the homeostatic behavior relied on adaptation of G1 duration as well as growth rate modulations. We developed a general mathematical framework that characterizes size homeostasis behaviors. Applying it on datasets ranging from bacteria to mammalian cells revealed that a near-adder is the most common type of size control, but only mammalian cells achieve it using modulation of both cell growth rate and cell-cycle progression.

## Introduction

While there is little consensus about the way mammalian cells control their size (Lloyd 2013; Ginzberg et al. 2015), studies of single-celled yeast and bacteria led to clearer picture. Cells achieve size homeostasis by adapting the amount of growth produced during a cell division cycle to their initial size, i.e. large cells should grow less, while small cells should grow more. There is a wide range of possible behaviors, but it is useful to exemplify size homeostasis by three simple limit cases: the sizer, the adder and the timer. Perfect size control was reported for the fission yeast, *S. Pombe* (Fantes 1977), where a size threshold (sizer) was proposed to control the passage of cells across several key cell cycle transitions (Pan et al. 2014; Wood & Nurse 2013). By contrast, an ‘adder’ mechanism relies on the addition of a constant volume at each cell cycle that is independent of initial size (Amir 2014; Voorn & Koppes 1998), causing cells to converge on an average size after a few generations. This behavior has been reported for several types of bacteria, cyanobacteria and in budding yeast (Campos et al. 2014; Taheri-Araghi et al. 2015; Soifer, Robert, Amir, et al. 2016; Yu et al. 2017; Deforet et al. 2015). Finally, if cells grow exponentially for a constant amount of time (a process called timer), large cells grow more than smaller ones and sizes rapidly diverge. Alternatively, if cells grow linearly, a timer results in cells growing by the same amount each cell cycle, therefore maintaining homeostasis (Conlon & Raff 2003). The growth pattern (i.e. exponential or linear) has therefore strong implications on the control needed to maintain size homeostasis.

In most cases, growth has been found to be exponential in bacteria and budding yeast (Godin et al. 2010; Wang et al. 2010; Osella et al. 2014; Iyer-Biswas et al. 2014; Di Talia et al. 2007; Soifer, Robert & Amir 2016) and bilinear in fission yeast (Horvath et al. 2016; Sveiczer et al. 1996; Mitchison 2003; Nobs & Maerkl 2014). Importantly, in all these organisms, size homeostasis has been reported to rely on an adaptation of cell cycle duration to initial size (reviewed in(Turner, Ewald, and Skotheim 2012; Osella et al. 2017; Jorgensen and Tyers 2004). The way such coordination between cell cycle progression and size is achieved is subject to intense debate. Indeed, the same effective behavior at the phenomenological level can result from a unique mechanism (Harris & Theriot 2016; Soifer, Robert & Amir 2016; Ho & Amir 2015) or from the combination of distinct mechanisms acting either in parallel (Osella et al. 2014) or sequentially through different cell-cycle sub-periods (Adiciptaningrum et al. 2015; Wallden et al. 2016). When several regulatory processes are involved, the overall emergent pattern is likely to be more complex than the stereotypical adder or sizer (Jun & Taheri-Araghi 2015; Sauls et al. 2016; Osella et al. 2017; Willis et al. 2016; Kennard et al. 2016).

The spectacular progress in understanding how unicellular organisms control their size was made possible thanks to high-throughput single live cell size tracking (Wang et al. 2010; Iyer-Biswas et al. 2014; Nobs & Maerkl 2014). However, similar progress has yet to be made in mammalian cells which have complex and fluctuating shapes.

As a result, most studies on mammalian cells have relied on population level measures (Conlon & Raff 2003; Dolznig et al. 2004; Echave et al. 2007; Killander & Zetterberg 1965; Kafri et al. 2013). These include attempts to extrapolate growth dynamics from size measurements at fixed time-points across a population (Kafri et al. 2013; Sung et al. 2013; Tzur et al. 2009). Recently, a variety of parameters have been used as proxies for size at the single cell level, mostly through indirect techniques (Popescu et al. 2014). These include cell dry mass (Park et al. 2010; Sung et al. 2013; Mir et al. 2011), buoyant cell mass (Son et al. 2012) and cell density (Grover et al. 2011). Among these recent studies, some have reported single live cell measurement of size at specific times in the cell cycle (Varsano et al. 2017) or through complete cell cycles (Mir et al. 2011; Park et al. 2010; Son et al. 2012). Although most data in unicellular organisms were obtained on volume, and most size-sensing mechanisms currently debated are thought to involve concentration-dependent processes (Ho & Amir, 2015; Schmoller et al., 2015; Sompayrac & Maaloe, 1973; Zielke et al., 2011), measurements of volume trajectories on single cycling mammalian cells have not been reported yet. This has limited the identification of regulatory processes leading to size homeostasis in mammalian cells.

Given the amount of evidence that unicellular organisms control their size by modulating their cell cycle progression, the same hypothesis was also made in mammalian cells. Similarly to budding yeast (Fisher 2016), a role for G1 duration was suggested (Dolznig et al. 2004; Killander & Zetterberg 1965; Conlon & Raff 1999) and verified directly for the first time very recently (Varsano et al. 2017). This last study showed that G1 duration could adapt as a function of birth size down to a minimum duration, leading, for these cells, to strong size-control for small-born cells and weaker, adder-like behavior for larger cells. Other studies on mammalian cells have reported negligible changes in cell cycle timing and have proposed that growth, although exponential on average, is modulated at specific points across the cell cycle and that changes in growth speed contribute to cell size control (Tzur et al. 2009; Kafri et al. 2013, for clarity, we recall here the definition of growth speed, which is the evolution of volume as function of time, and of growth rate, which is the evolution of growth speed as a function of volume). Direct observation of a convergence of growth speed at G1/S transition was provided for lymphoblastoid cells (Son et al. 2012) but how this could lead to an effective homeostatic behavior was not characterized. The idea that growth speed modulations could play a role in mammalian cells size control was never tested directly, nor has its relative contribution to the effective homeostatic behavior been compared to that of time modulation. Moreover, it is not known how the two-tier mechanism found recently in G1 phase (Varsano et al. 2017) extends to cells growing exponentially or whether it generalizes to other cell types.

To address these questions as directly as possible, we recently developed two new methods to precisely measure the volume of single live cells over several days, on a large number of cells, independently of their shape (Cadart et al. 2017; Zlotek-Zlotkiewicz et al. 2015; C. Cadart et al. 2014). In this study, we used these tools to track single-cell volume growth over complete cell cycles. To understand how the coupling of growth and time modulations results in size control, we also developed a quantitative framework that characterizes the relative contributions of timing and growth modulations to size homeostasis from bacteria to mammalian cells.

## Results

### Single cell volume measurement over complete cell division cycles

It is a general belief that proliferating cultured cells double their size between two divisions, yet the relation, for single cells, between their size at mitotic entry and their size at birth, has never been reported for freely growing mammalian cells in culture.

To establish this relation, it is necessary to track single proliferating cells and measure the volume of the same cell at birth and at mitotic entry. We implemented two distinct methods to obtain these measures. First, we grew cells inside microchannels of a well-defined cross-sectional area (Figure S1A, and (Cadart et al. 2014)). In such a geometry, dividing cells occupied the whole section of the channels and we could infer their volume from their length, like for yeasts and bacteria. The second method we used was based on fluorescence exclusion (FXm, (Zlotek-Zlotkiewicz et al. 2015; Cadart et al. 2017), Figure 1A, Movie S1) and has been optimized for long term recording and automated analysis of populations of growing cells ((Cadart et al. 2017), Figure S1B and supplementary information). This method has several advantages compared with the channels: it does not require growing cells in a very confined environment which is thought to constrain growth to a linear pattern (Varsano et al. 2017), it is more precise and also produces complete growth trajectories for single cells (Figure 1B-C and S1C). Visual inspection of the movies was used to determine key points in the cell division cycle for each single cell tracked. Volume at birth was defined as the volume of a daughter cell 40 minutes after cytokinesis onset, while volume at mitotic entry was defined as volume of the same cell 60 minutes prior to the next cytokinesis onset (Figure 1B). These intervals were chosen to avoid the period of volume overshoot corresponding to mitosis (Zlotek-Zlotkiewicz et al. 2015; Son et al. 2015) (Figure S1D-E). Analysis of growth speed as a function of size, for a large number of single cells and cell aggregates showed global exponential growth, as expected for freely growing cells (Figure S1F) (Tzur et al. 2009; Mir et al. 2011; Sung et al. 2013; Son et al. 2012). For each experiment performed, the dataset was checked for quality: we verified that the distribution of volumes at birth and the average growth speed did not change throughout the experiment, and that these values did not change from one experiment to another (Figures 1D and S1G). All the experiments in our dataset match these quality criteria (note that we kept one dataset which showed a significant, but small, decrease in volume through the course of the experiment, because despite optimization, we could not avoid some internalization of dextran by these cells, Figure S1G, HeLa cells). We were thus able, with these methods, to produce fully validated high quality datasets of single cell volume along cycles of growth and division, which can be further used to ask elementary questions on volume homeostasis for proliferating cultured mammalian cells.

**Figure 1:**
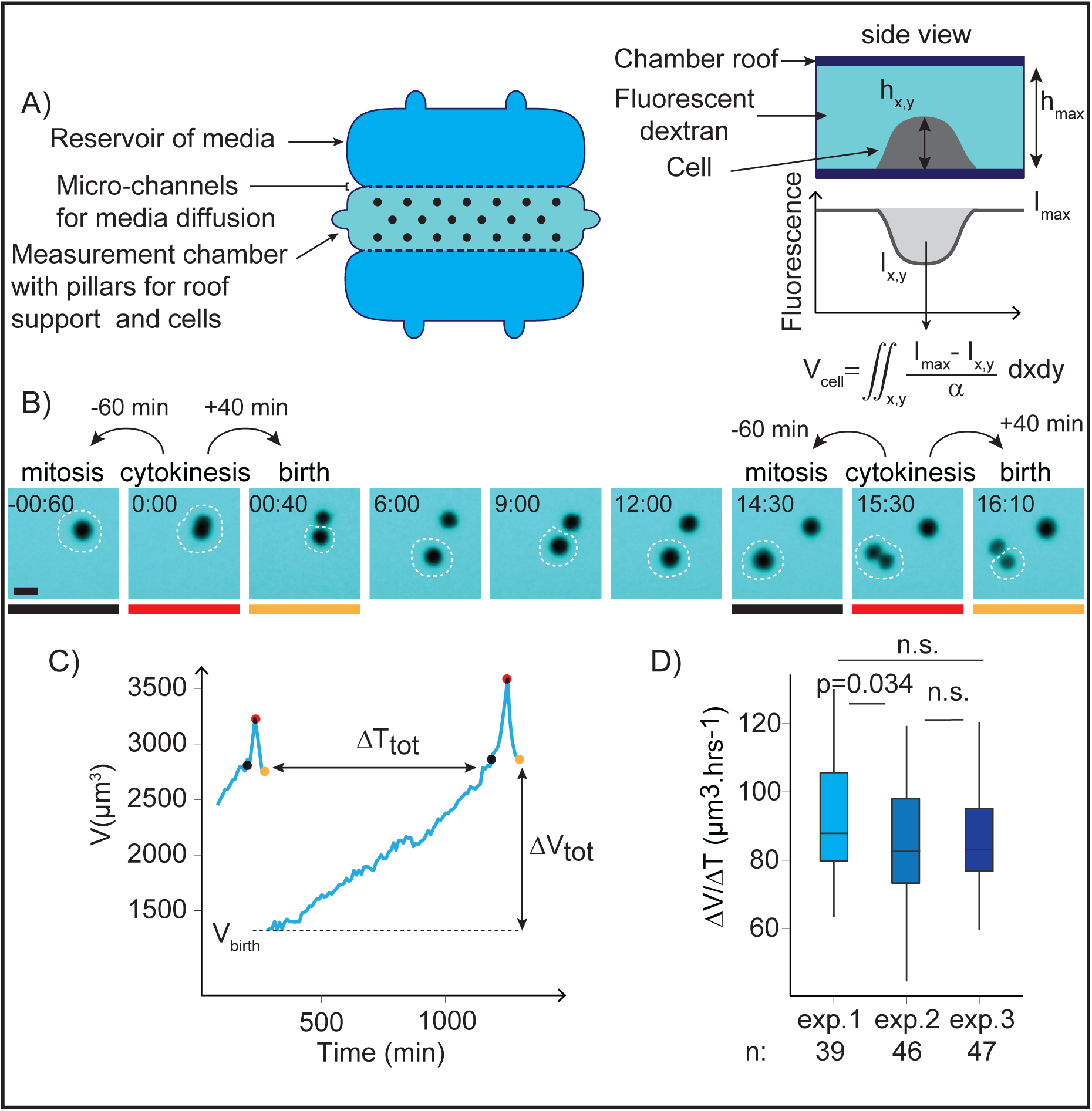
Single-cell volume tracking over entire cell division cycles. A) Principle of the fluorescence exclusion-based volume measurement method (FXm). Left: measurement chamber used for 50hrs long time-lapse acquisitions (see SI). Right: principle of the measure. Fluorescence intensity at a point *γ*_*x*,*y*_ of the cell is proportional to the height of the chamber minus the height *h*_*x*,*y*_ of the cell at this point. Fluorescence intensity *γ*_*max*_is the intensity under the known height of the chamber roof *h*_*max*_, where no object excludes the fluorescence. Integration of fluorescence intensity over the cell area gives the cell volume *V*_*cell*_ after calibrating the fluorescence intensity signal *α* = (*I*_*max*_ − *I*_*min*_) / *h*_*max*_ (see SI). B) Sequential images of a HT29 cell acquired for FXm. Mitosis and birth are defined as the time-points 60 min before and 40 min after cytokinesis respectively. The white dashed circle indicates the cell measured in C), the colored lines indicate the time-points highlighted by circles of the same color in C). Time is in hours:minutes. Scale bar is 20 μm. C) Single HT29 cell growth trajectory (volume as a function of time) and key measurement points (see SI). The time-points shown in B) and underlined in grey, red or yellow are indicated by points of matching colors on the curve in C): the grey points correspond to volume at mitotic entry, the red points correspond to volume at cytokinesis and the yellow points to volume at birth. ΔT_tot_ is the total duration of the cell division cycle from birth to mitosis and ΔV_tot_ is the total added volume. D) Average growth speed for three independent experiments with HT29 wild-type cells. n= 39 (exp. 1), n=46 (exp. 2), n=47 (exp. 3). The hinges show the 25th and 75th percentiles and the bars extends from the hinge to the highest value that is within 1.5 * IQR (Inter Quantile Range) of the hinge. The p values are the result of a pairwise t test comparing the means. See also Figure S1 and Movie S1

### A near-adder behavior is observed in cultured mammalian cells

The first elementary information comes from the relation between the added volume and the volume at birth. We thus made that plot, together with the equivalent plot of volume at mitotic onset versus volume at birth, for each cell line and condition in our dataset (Figure 2A and S2A). If cells were doubling their volume (i.e. in the case of exponentially growing cells with a timer), the added volume would be equal to the volume at birth, thus the two values would linearly correlate with a slope of 1, and the final versus initial volume plot would show a slope of 2. On the other hand, if cells were perfectly correcting for differences in size, added volume would be smaller for bigger cells, so the slope would be negative, while the final volume would be identical for all cells independently of their initial volume. We studied 3 different cancerous cell lines (HT29, HeLa, and Raji) and on Madin-Darby canine kidney (MDCK) epithelial cells and found neither of these two extremes. With the exception of Raji cells (human B lymphoblast), which showed a large dispersion of added volumes, and for which added volume slightly correlated with initial volume (Figure S2A), we instead found that added volume showed no correlation with initial volume (Figure 2A, left panel). Consistently, the volume at mitotic entry showed a clear linear correlation with volume at birth, with a slope close to 1 (Figure 2A, right, and Figures 2B and S2A), suggesting that all cell lines studied, except the Raji cells, grew by the same amount, on average, independently of their volume at birth. This observation was also reproduced when analyzing previously published results obtained on lymphoblastoid L1210 cells (kindly shared by the authors (Son et al. 2012)). This behavior is close to that of an adder, and was already described for several bacterial species and for the buds of budding yeast cells (Campos et al. 2014; Taheri-Araghi et al. 2015; Soifer, Robert, Amir, et al. 2016). This weak form of volume homeostasis was shown, theoretically and experimentally for bacteria and yeasts, to be able to compensate for asymmetries in sizes during division. A direct prediction is that, after an asymmetric division, the difference in size of the two daughter cells would be reduced by half in one division cycle, but not completely compensated. We artificially induced asymmetric divisions by growing cells inside microchannels (Figure S1A, Movie S2). Confinement prevents mitotic rounding, which leads to errors in the mitotic spindle positioning and ultimately generates uneven division of the mother cell (Figures 2C-D, (Lancaster et al. 2013; Cadart et al. 2014)). We then compared the asymmetry in volume, at birth and at the next mitosis, between pairs of daughter cells that had divided inside these channels. We found that their level of volume asymmetry at birth was much higher than in control cells that divided outside of the channels, and that it was significantly reduced at entry into the next mitosis, but not completely compensated (Figure 2D), as predicted with an adder behavior. In conclusion, this first analysis of our dataset revealed that most cultured mammalian cell lines display a near adder behavior. Importantly, a near-adder observed at the phenomenological level does not necessarily imply the existence of a molecular mechanism ‘counting’ added volume. The most recent findings in budding yeast and *E. coli* rather propose that the adder emerges from the combination of several mechanisms acting in parallel or sequentially during the cell cycle (Osella et al. 2017).

**Figure 2:**
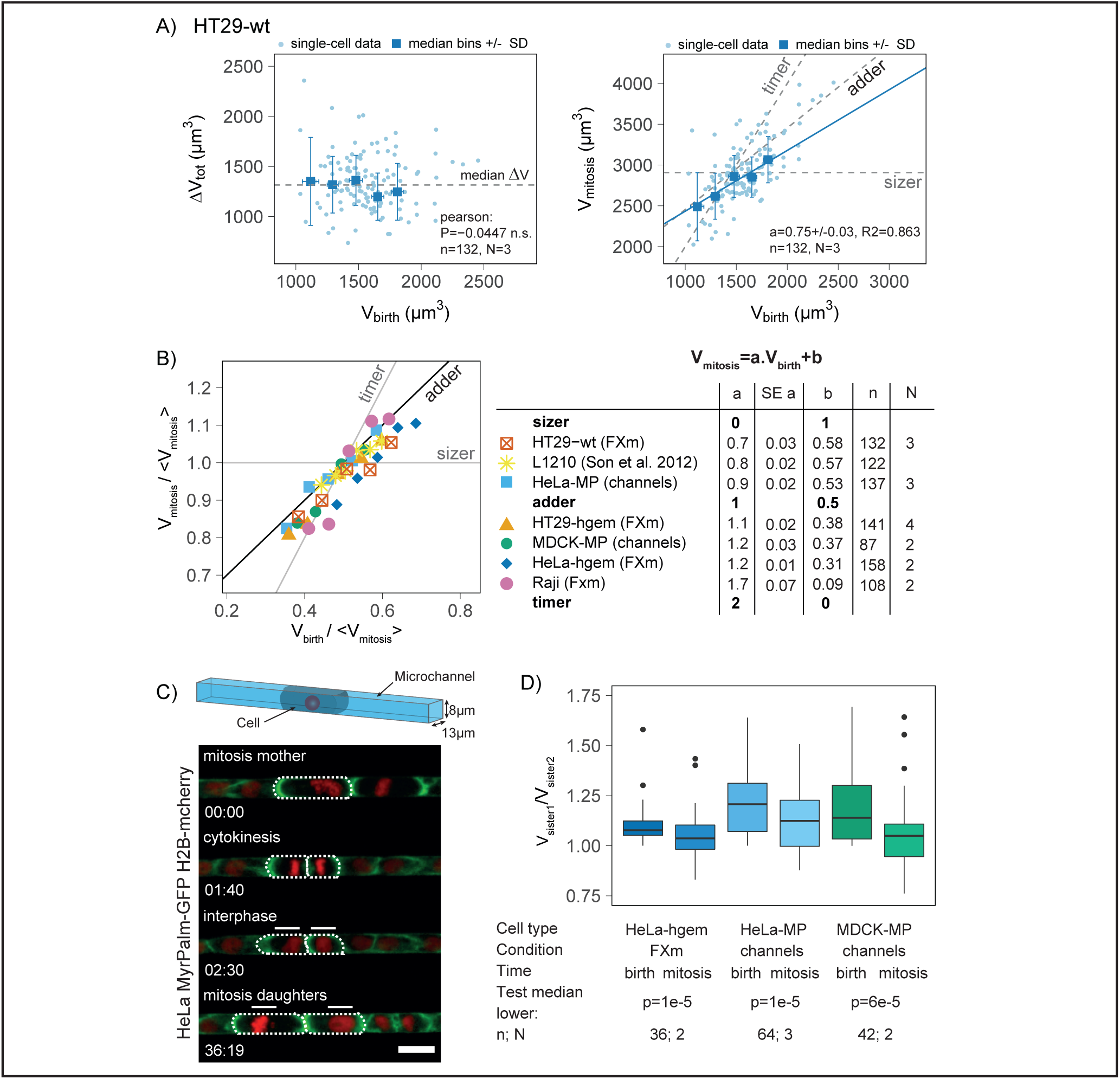
Adder-like behavior in cultured mammalian cells. A) Left: total volume gained during one cell division cycle ΔV_tot_ versus volume at birth V_birth_ for wild-type HT29 cells. The dashed grey line shows median added volume. Right: volume at mitotic entry V_mitosis_ versus volume at birth V_birth_. Dashed grey lines show the expected trends in case of a sizer (horizontal line at average volume at mitosis), an adder (line of slope 1 with an intercept at mean volume at birth) and a timer (line of slope 2 with intercept at zero). Blue line: linear fit on the binned data weighted by the number of observations in each bin (see SI), with a the slope of the linear fit +/− standard error and R2 is the coefficient of determination, n is the number of single-cell observations and N the number of independent experiments. B) Left graph: comparison of volume at mitosis rescaled by the mean volume at mitosis versus volume at birth rescaled by the mean volume at mitosis for various cultured mammalian cell lines. Ideal curves for stereotypical homeostatic behaviors are shown as black lines. The points corresponds to median bins (the graphs with single cell points and median bins for each cell type are shown in Figure S2A). For each cell type, a linear fit V_mitosis_ = a. V_birth_ + b is made on the bins weighted by the number of observation in each bin (see SI). Right table: estimates from the linear regression for each cell type: a (slope coefficient), SE a (standard error for a), b (slope intercept), n (number of single cell observations) and N (number of independent experiments). The theoretical slope coefficients and intercepts expected in case of sizer, adder or timer are also indicated. The HT29 are human colon cancer epithelial cells (wild-type (HT29-wt) or expressing hgeminin-mcherry (HT29-hgem)), HeLa are human cervix epithelial cancer cells (either stably expressing MyrPalm-GFP and Histon2B-mcherry (HeLa-MP) or hgeminin-GFP (HeLa-hgem)), MDCK expressing MyrPalm-GFP (MDCK-MP) are madine canine kidney epithelial cells, Raji are human B lymphoblastoid cells, L1210 are mouse lymphoblastoid cells from data kindly sent by Son and colleagues (Son et al. 2012). Apart from the L1210 cells buoyant mass, data are volumes acquired with either the FXm or the microchannel methods (See C)). C) Top: scheme of a cell confined in a microchannel (nucleus in red). Bottom: sequential images of an asymmetrically dividing HeLa cells expressing MyrPalm-GFP (plasma membrane, green) and Histon2B-mcherry (nucleus, red) growing inside a microchannel. The outlines of the cell of interest and its daughters (white bars) are shown with white dotted lines. Scale bar is 20 μm. Time is hours:minutes. D) Ratio of volume in pairs of sister cells at birth and mitosis for MDCK cells expressing MyrPalm-GFP (MDCK-MP) and HeLa cells expressing MyPalm-GFP and Histon2B-mcherry growing inside microchannels (HeLa-MP). Control, in non-confined condition, corresponds to HeLa expressing hgeminin-GFP (HeLa-hgem) cells measured with FXm. A Wilcoxon signed rank test was performed to test that the median ratio was lower from birth to mitosis in each condition. The upper and lower hinges of the boxplot represent the 25th and 75th percentile, the bars extend from the hinge to the highest (lowest) value within 1.5*IQR (Inter Quantile Range) of the hinge. Data beyond the whiskers are shown as outliers. n indicates the number of pairs of sister cells and N the number of independent experiments. See also Figure S2 and Movie S2

### G1 duration is negatively correlated with volume at birth

Modulations of cell cycle duration as a function of size are the core of size regulation in unicellular organisms. Such modulations were therefore the natural hypothesis to explain size control in mammalian cells and an adaptation of G1 duration to initial size, very similarly to what had been shown in budding yeast (Hartwell & Unger 1977; Johnston et al. 1977) was proposed (Killander & Zetterberg 1965; Dolznig et al. 2004). Recently, direct evidence supporting this was provided (Varsano et al. 2017) while other contributions argued against the existence of time modulation (Son et al. 2012; Kafri et al. 2013), opening a controversy on this question, with few direct observation available to clarify this point. In our dataset, cells grown in the volume measurement chamber, but not inside microchannels, showed a longer cell division cycle for smaller cells (Figure S2A, middle graphs). To investigate this point in more details, we combined cell volume measurements on HT29 cells with a classical marker of cell cycle phases, hgeminin-mcherry, which accumulates in the cell nucleus at S-phase entry (Sakaue-Sawano et al. 2008) (Figures 3A, S3A and Movie S3). This new dataset confirmed that, at the scale of the entire cell division cycle, cells added the same amount of volume independently of their volume at birth. During G1 phase, small cells at birth added slightly more volume than large ones, while during S-G2, large cells at G1/S added slightly more volume than small ones (Figure 3B and S3B, left graphs). Consistently, the volume at G1/S transition plotted against volume at birth showed a slope below 1 (a = 0.7 ± 0.01), suggesting a homeostasis mechanism more efficient than an adder, while the slope of the volume at mitosis entry versus volume at the G1/S transition was 1.4 ± 0.02, suggesting a poor homeostasis mechanism (Figure S3B, right graphs). Consistent with a regulation occurring mostly in G1, the distribution of G1 durations was wide and right-skewed, resembling the distribution of entire cell division cycle durations (Figures 3C and S3C, CV=53%), while S-G2 showed a very narrow and symmetrical distribution of durations (Figures 3C and S3C, CV=18%) (Fligner-Kileen test comparing the standard deviations, p=2*10^-16). Consistently, the duration of G1 was highly correlated with the total duration of the cell cycle, while it was less for S-G2 (Figure S3D). Plotting the time spent for single cells in a given phase versus its volume at the beginning of that phase, confirmed that smaller cells at birth had a longer G1 phase (Figure 3D). By contrast, S-G2 duration was not correlated with volume at entry in S phase (Figure S3B, middle graphs). This analysis also suggested that there was a minimal time cells spent in G1, and that dispersion of G1 duration was larger for smaller than for larger cells, which tended to spend only a minimal time in G1 (about 4 to 5 hours) (Figure 3D). This is well illustrated by the cumulative distribution function of the time spent in G1 for various ranges of volumes at birth (Figure 3E). We could not distinguish between the effective adder observed from birth to mitosis and the alternative model currently debated in yeast (Soifer, Robert, Amir, et al. 2016) and bacteria (Ho & Amir 2015) where the constant volume is added between two replication initiation events (see supplementary information, Figures S3E-G). These data together suggest that, despite an overall exponential growth, smaller cells can add, on average, as much volume as bigger cells, thus achieving an adder behavior, by extending the duration of the G1 phase, while S-G2 phase rather resembles a timer, with a duration independent of size at the G1/S transition.

**Figure 3:**
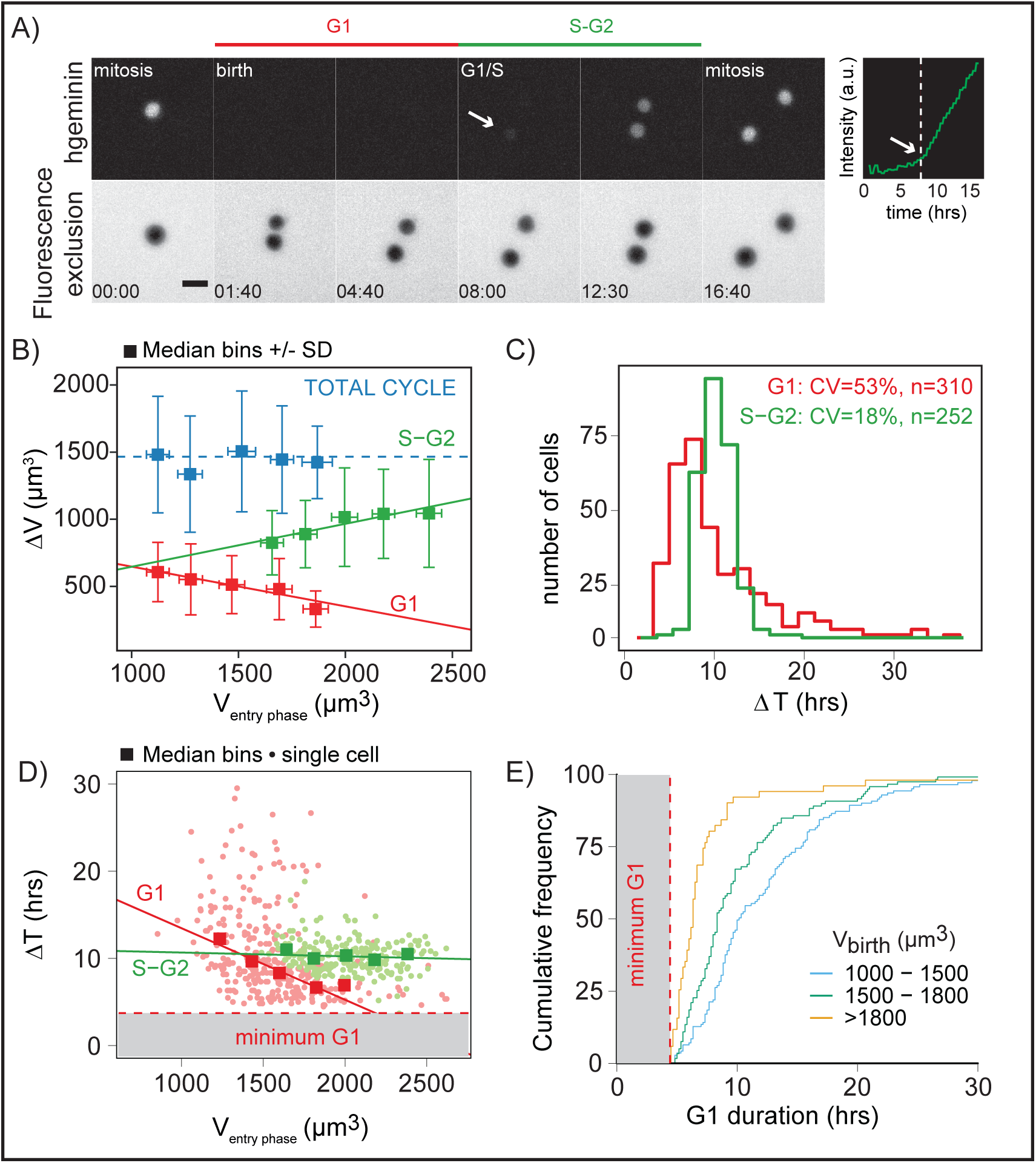
For HT29 cells, G1 duration is anticorrelated to V_birth. A) Sequential images of HT29 cells expressing hgeminin-mcherry (top) and FXm on the same cells (bottom). The graph on the right shows the quantification of hgeminin-mcherry in the cell as a function of time. Time zero corresponds to mitosis (see SI). The vertical white line and arrows indicate the time at which hgeminin-mcherry becomes detectable. G1 phase (red line) spans from birth to appearance of hgeminin (G1/S transition) and S-G2 phases (green line) from G1/S to next entry in mitosis (see SI). Scale bar is 20μm. Time is in hours:minutes. B) Total added volume ΔV in G1 (red), S-G2 (green) and the total cell cycle (blue) as a function of volume at entry in the phase (volume at birth, volume at G1/S and volume at birth respectively). Dashed blue line indicates the median volume added during the total cell cycle. Squares show median bins with standard deviation (bars). Pearson’s correlation coefficient (P), for the entire cell division cycle: P=0.13, p=0.12, n=141; G1: P=−0.30, p=9*10^-7, n=241; S-G2: P=0.34, p=1*10^-5, n=144; linear regression on the median bins weighted by the number of observations in each bin, G1: a= −0.30+/−0.009, p=1*10^−91, R2=0.83; S-G2: a=0.3+/−0.01, p=3*10^−53, R2=0.83. C) Histogram of G1 phase, ΔT_G1_ (red) and S-G2 phases ΔT_S-G2_ (green). CV: coefficient of variation. Standard deviations are significantly different (Fligner-Killeen test, p=2*10^-16). D) Duration of G1 phase, ΔT_G1_, and S-G2 phases, ΔT_S-G2_ versus volume at entry in the phase (V_birth_ and V_*G*1/*S*_ respectively). Individual cell measures (dots) as well as median bins (squares) are shown. Pearson’s correlation coefficient, G1: P=−0.33, p=7*10^-10 (n= 310); S-G2: P=−0.12, p=0.07 (n=220); linear regression on the median bins weighted by the number of observations in each bin, a +/−standard error (slope coefficient), p value and coefficient of determination (R2) for G1: a=−0.008+/−0.0001, p=1*10^-160, R2=0.92; for S-G2: a=−0.004+/−0.0001, p=9*10^-4, R2=0.07. Red dashed line and grey area are a visual guide for minimum G1 duration around 4-5 hours. E) Cumulative frequency graph of ΔT_G1_ binned for three ranges of volumes at birth V_birth_. Red dashed line and grey area are a visual guide for minimum G1 duration around 4-5 hours. All data in the figure are from HT29 cells expressing hgeminin-mcherry (N=4). See also Figure S3 and Movie S3

### Abnormal large cells do not adapt G1 duration but modulate their growth rate

Figures 3D-E show a lower limit on the duration of G1 phase, which implies that, if growth was exponential and homeostasis limited to the G1 phase, it would not be possible to have homeostasis for larger cells. To produce larger cells at birth, we arrested cells using Roscovitine, an inhibitor of major interphase cyclin dependent kinases, like Cdk2 (Meijer & Raymond 2003). For this experiment, we used HeLa cells, because HT29 cells, despite long arrest with Roscovitine treatment, only slightly increased their volume. After a 48hours block with Roscovitine, the drug was rinsed, and cells were injected in the volume measurement chamber. Recording started after the first mitosis following the release from Roscovitine. As expected, cells which had been treated with Roscovitine were on average 1.7 fold larger than the controls (Figure 4A and 4B, top histogram of the graph, Figure S4A). Single cell growth curves showed that Roscovitine treated cells behaved similarly to large control cells (Figure 4A, Movie S4). As expected, their G1 duration was shorter (Figure 4B right axis) and was on average closer to a minimal G1 duration (≈ 4 hours) independently of volume at birth (Figure 4B), a behaviour reminiscent of the recent results from (Varsano et al. 2017). Interestingly, this duration was the same as the duration displayed by large control cells. But, surprisingly, large Roscovitine treated cells still grew, during G1, by nearly the same amount of volume than smaller control cells, independently of volume at birth (Figure 4C). In S-G2 however, the duration was longer for Roscovitine-treated cells, maybe due to replication defects, and the added volume was thus also larger (Figure S4B-C). If G1 duration is not modulated and larger cells grow by the same amount, an alternative mechanism is that growth speed is modulated. We thus analyzed single cells growth curves in G1 and plotted their growth speed against their volume. It showed that, both for control cells and Roscovitine treated cells, and for all the range of sizes, growth speed in G1 increased with size, compatible with an exponential growth even for the largest cells (Figure 4D for G1, S4D-E for S-G2 and complete cell cycle, and S4F relative to G1/S transition). To understand how larger cells at birth could grow by approximately the same amount as smaller cells, in a similar amount of time, and because single growth curve trajectories showed complex behaviors (Figure S4G-I), we grouped Roscovitine and control cells and binned them by their size at birth. We then plotted, for small, intermediate and large cells at birth, their growth speed versus their volume. By definition, the slope of such plot indicates the growth rate. This showed that although, for all ranges of size at birth, growth was compatible with exponential, the slope of growth speed versus volume decreased for larger sizes at birth, suggesting a lower exponential growth rate for cells born bigger (Figure 4E). In conclusion, consistent with a minimal G1 duration, Roscovitine treated cells, which were as big or larger at birth than large control cells, displayed a G1 duration independent of their volume at birth and on average equal to the minimal G1 duration found for control cells. They still added the same amount of volume as smaller cells, due to a modulation of their exponential growth rate, with larger cells at birth showing a smaller growth rate. By producing larger cells at birth, we were able to reduce cell cycle modulation and reveal the existence of a strong growth rate modulation.

**Figure 4:**
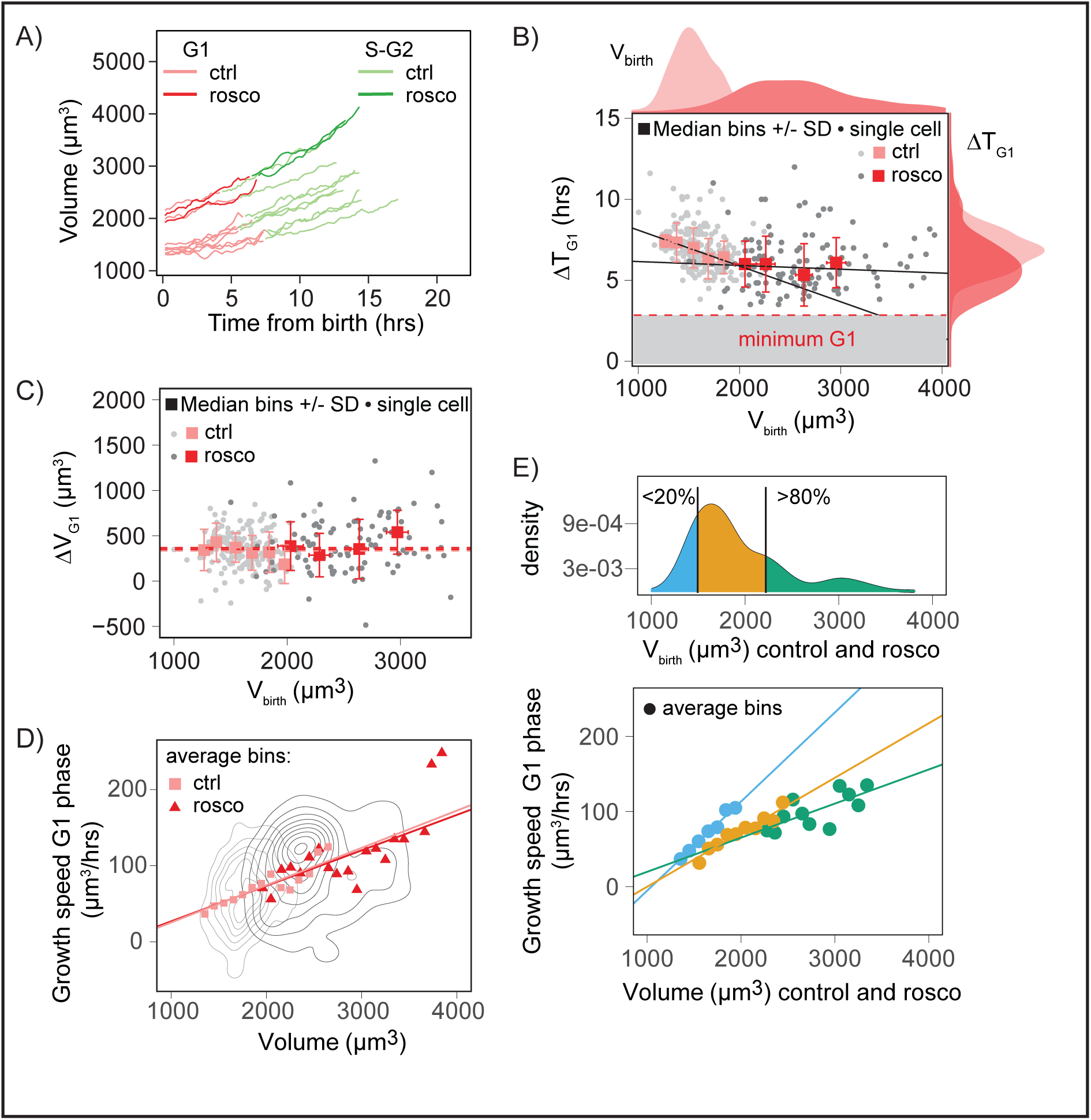
Larger cells obtained after Roscovitine treatment still display an adder behavior. A) Examples of single-cell growth trajectories for HeLa cells expressing hgeminin-GFP, either control (“ctrl”, lighter color), or after washout from Roscovitine treatment (“rosco”, darker color) as a function of time from birth; B) G1 is in red and S-G2 in green. More examples can be found in Figure S4H-I. G1 duration, ΔT_G1_, as a function of volume at birth for control (“ctrl”, light red, light grey) and Roscovitine-treated cells (“rosco”,darker red, darker grey). Individual cell measures (dots) as well as median bins (squares) and standard deviation (bars) are shown. Red lines shows linear regression on median bins weighted by the number of event in each bin (a (slope coefficient) +/− standard error, R2 (coefficient of determination) and p value; control: a=−0.00188+/0.00008, R2=0.76, p=5*10-61; Roscovitine: a=−0.0002±0.0001, R2=0.05, p=1*10-42; Pearson’s correlation coefficient, control: P=−0.4, p=3*10^-8, n=199, N=2; Roscovitine: P= −0.03, p=0.8., n=120, N=3). Note that control HeLa cells are already very close to the minimum G1 duration compared to the HT29 cells shown in Figure 3. Distributions on the top of the graph are the distribution of volume at birth V_birth_; control: mean volume at birth=1600μm3, n=231; Roscovitine: mean volume at birth=2600μm3, n=136; Welch t test comparing the means: p=2.2*10^-16. Distributions on the right of the graph show G1 duration; control: mean G1 duration=7.0hrs, n=201, N=2; Roscovitine: mean G1 duration=6.1hrs, n=124, N=3; Welch t test comparing the means: p=6.5*10^-7. Red dashed line and grey area are a visual guide for minimum G1 duration around 4-5 hours. C) Added volume in G1 (ΔV_G1_) versus volume at birth for control (“ctrl”, light red, light grey) and Roscovitine-treated cells (“rosco”, darker red, darker grey). Individual cell measures (dots) as well as median bins (squares) and standard deviation (bars) are shown. Pearson’s correlation coefficient, control: P=−0.06, p=0.4, n=178, N=2; Roscovitine: P=−0.07, p=0.8, n=108, N=3. Dashed lines represent the mean added volume in each condition, control: mean added volume in G1=350 μm3; Roscovitine: mean added volume in G1=390 μm3; Welch’s t.test comparing the means, p=0.2423. D) Instantaneous growth speed δv/δt in G1 as a function of volume, for control (“ctrl”, light red squares, number of cells n=119, N=1) and Roscovitine treated cells (“rosco”, dark red triangles, number of cells n=49, N=2). Growth speed was estimated on sliding windows of 90 min (see Figure S4G-I and SI). Red lines shows linear regression on average bins weighted by the number of event in each bin (slope coefficient a +/− standard error, p value and R2 (coefficient of determination) are: for the control cells: a=0.0489+/−0.0005, p≈0, R2=0.78; for the Roscovitine treated cells: a=0.047+/−0.002, p=1*10^-137, R2=0.49). Grey lines represent bivariate kernel densities. E) Top: kernel density of volume at birth for control and Roscovitine treated cells together. The 20% and 80% percentiles are represented. Bottom: Same data as D) but cells are grouped by their initial volume at birth (cells within the 0 to 20% percentile (blue), 20 to 80% percentile (orange) and 80 to 100% percentile of the distribution (green). The dots represent the averaged bins which contain measurements on at least 5 different cells, the lines are robust linear fits on the bins. For each group, the values of a (slope coefficient of the fit) +/− standard error, p value of a, R2 (coefficient of determination), nc (number of cells from the control condition in the group) and nr (number of cells from the Roscovitine condition in the group) are: 0-20%: a=0.119+/−0.008, p=4.1*10^-5, R2=0.98, nc=24, nr=0; 20%-80%: a=0.072+/−0.009, p=4.88*10^-5, R2=0.90, nc=60, nr=15; 80%-100%: a=0.05+/− 0.01 p=0.00192, R2=0.43, nc=3, nr=24; N number of independent experiments is control: N=1, rosco: N=2. All data are from HeLa cells expressing hgeminin-GFP. See also Figure S4 and Movie S4

### A general mathematical framework to compare the homeostatic process from bacteria to eukaryotes

Our results show evidence of time modulation in G1, in agreement with recent findings (Varsano et al. 2017) and directly support the hypothesis that modulations of growth rate might also contribute to size homeostasis (Kafri et al. 2013). To understand the respective contribution of growth and time modulation on the effective homeostatic process, we built a general mathematical framework. Such contribution of growth rate modulation has never been reported in unicellular organisms and our framework allowed us to perform a comparative analysis of the way by which mammalian cells and unicellular organisms achieve size homeostasis. Our model (described in details in the supplementary information) is applicable to the whole cell cycle or to single cell-cycle stages (for the sake of simplicity, we will discuss it hereon for an entire cycle). It assumes that cells grow exponentially, which corresponds to the most common behavior we observed in our dataset, and adopt a rate chosen stochastically from a probability distribution. This rate may depend on volume at birth (and hence contribute to size correction). Similarly, the interdivision time (cell cycle duration) may be chosen based on volume at birth and has a stochastic component. Correlations between growth rate, interdivision time and size at birth are accounted to linear order, motivated by the fact that such linear correlations are able to explain most patterns in existing data (at least for bacteria, (Grilli et al. 2017)). The resulting model is able to characterize the joint correction of size by timing and growth rate modulation, with a small number of parameters.

A first parameter, λ, describes how the total relative growth (log(*V*_*mitosis*_/*V*_*birth*_)) depends on volume at birth. If λ = 1, the system behaves like a sizer, if it is 0.5, it is an adder and if it is 0, there is no size control at all (on average, cells divide when they doubled their initial volume). This parameter can be described, for each dataset, by performing a linear regression on the plot of log(*V*_*mitosis*_/*V*_*birth*_) versus the log(*V*_*birth*_) (Figures 5A, S5A and equation 5 in supplementary informations). The second parameter, *θ*, describes how interdivision time depends on volume at birth. This parameter can be described, for each dataset, by performing a linear regression on the plot of cell cycle duration (τ = Δ*T*) versus log(*V*_*birth*_) (Figures 5B, S5B and equation 6 in supplementary informations). If this correlation is negative (which, by choice, corresponds to a positive value of the parameter meant to describe the strength of the correction), it means that larger cells will tend to divide in shorter times, hence these cells operate size correction due to a modulation of timing. Finally, the third parameter, *γ*, describes the link between initial size and a variation in growth rate with respect to its mean value. Similarly, if *γ* is positive, modulations of growth rate positively contribute to size control. This can be obtained by linear regression when the corresponding measurements are available (e.g. in data from bacteria, (Wallden et al. 2016; Kennard et al. 2016; Kiviet et al. 2014; Taheri-Araghi et al. 2015) Figures 5C and S5C, equation 4 in supplementary information). When growth rate for single-cells was not available (for mammalian cells and yeasts), the parameter *γ* was estimated from the values of λ and *θ* using an approach based on the covariance between pairs of measured variables (Figure S5D-E and supplementary information). The validity of this approach was tested on the bacteria datasets where all parameters can be directly measured (Figure S5E and supplementary information).

These three parameters are linked by a balance relation, which describes the fact that the overall size correction results from the combination of timing and growth rate corrections (see also supplementary information).

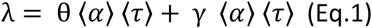

Each cell line and condition can be characterized by one value for each parameter and thus one point on the graph which shows *γ* versus *θ* (Figure 5D). Additional (less relevant here) parameters concern the intrinsic stochasticity of interdivision timing, growth rates and net growth (see supplementary information). For eukaryotes where the growth rate ⟨α⟩ is not easily accessible, the product ⟨*α*⟩ ⟨τ⟩ was approximated by: ⟨*α*⟩ ⟨τ⟩ ≈ ⟨*G*⟩ = ⟨log(*V*_*mitosis*_)/log(*V*_*birth*_)⟩ (Figure 5D, right, supplementary information). The validity of this normalization was tested with bacteria (Figure S5F and supplementary information).

**Figure 5:**
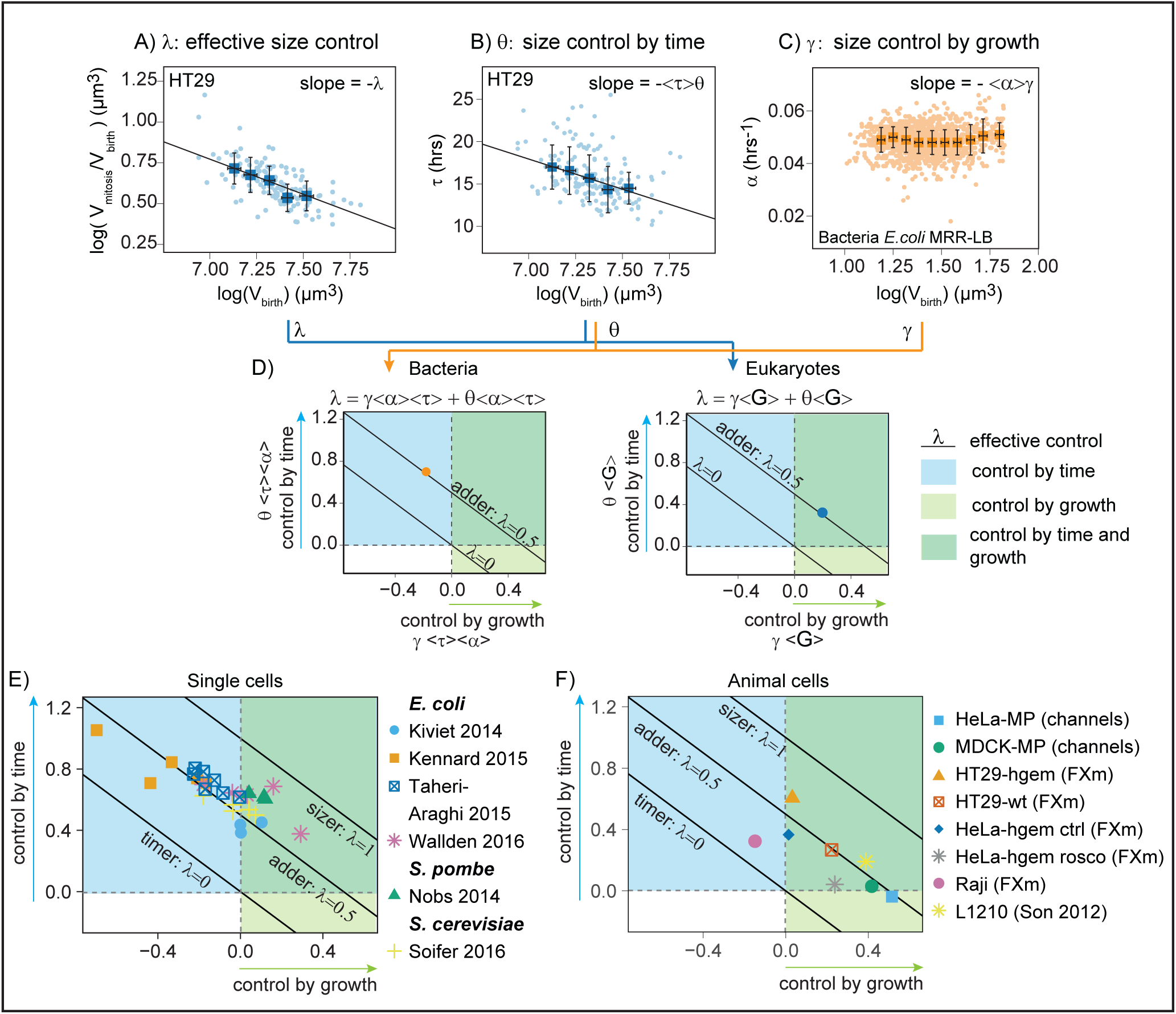
Contribution of growth and time modulation in overall size control. A) Doubling rate, log(V_mitosis_/V_birth_) versus logarithm of initial volume log(V_birth_) for HT29 wild-type cells. The slope coefficient of the linear regression (performed on the median bins, blue squares, and weighted by the number of observations in each bin) gives -λ and indicates the effective size control, (-λ =−0.5±0.002, R2=0.85, n=132, N=3). Error bars represent standard deviation. Dots are single-cells, squares with error bars are median bins with standard deviation, the black line shows the linear regression. B) Cell cycle duration τ versus initial volume log(V_birth_) for HT29 wild-type cells. The slope coefficient of the linear regression (performed on the median bins, blue squares, and weighted by the number of observations in each bin) gives, by definition in our equations (see SI, Equation 6), -⟨τ⟩ *θ* where ⟨τ⟩ is the average cell cycle duration, and *θ* is the strength of control by time modulations. A positive value of *θ* corresponds to a positive effect on size control (-⟨τ⟩*θ* =−7±0.2, R2=0.88, n=163, N=3). Dots are single-cells, squares with error bars are median bins with standard deviation, the black line shows the linear regression. C) Growth rate a versus initial size log(V_birth_), for datasets on bacteria, from (Kennard et al. 2016). The slope coefficient of the linear regression (performed on the median bins, orange squares, and weighted by the number of observations in each bin) gives, by definition in our equations (see SI, Equation 4), -⟨*α*⟩*γ* the control due to growth rate modulations. A positive value of *γ* corresponds to a positive effect on size control (-⟨*α*⟩*γ* =−0.0005±0.0002, R2=0.06, n=2107). Dots are single-cells, squares with error bars are median bins with standard deviation. D) Left: plot of *θ* multiplied by the average cell cycle duration ⟨τ⟩ and the average growth rate ⟨*α*⟩, versus *γ* multiplied by ⟨τ⟩ and ⟨*α*⟩ for the bacteria dataset shown in C). Positive values along both y and x axes correspond to a positive effect on size control via time or growth modulation respectively. Right: plot of *θ* multiplied by ⟨*G*⟩, the average growth (⟨*G*⟩ = ⟨log(*V*_*mitosis*_)/log(*V*_*birth*_)⟩, versus *γ* multiplied by ⟨*G*⟩ for HT29 wild-type cells shown in A-B). The dashed lines indicate the threshold above which time modulation (horizontal line) and growth modulation (vertical line) have a positive effect on size control. E) Comparison of datasets for bacteria (data from (Wallden et al. 2016; Kennard et al. 2016; Taheri-Araghi et al. 2015; Kiviet et al. 2014)) and yeasts (data from (Soifer et al. 2016; Nobs and Maerkl 2014)), plotted as in D. The dashed lines indicate the threshold above which time modulation (horizontal line) and growth modulation (vertical line) have a positive effect on size control. Each point corresponds to a different growth condition (see Figure S5G). F) Comparison of datasets for animal cells (our results and data from (Son et al. 2012)), plotted as in D. The dashed lines indicate the threshold above which time modulation (horizontal line) and growth modulation (vertical line) have a positive effect on size control. See also Figure S5

Using these dimensionless parameters, it was then possible to compare datasets obtained from different cell types in different conditions and estimate whether they displayed volume homeostasis (λ > 0) with an adder behavior (λ = 0.5) or better (λ = 1). It was also possible to know if homeostasis relied more on time modulation (*θ* > 0) or growth rate modulation (*γ* > 0). With this framework (Figures 5E and S5G), all the datasets for both bacteria (Kennard et al. 2016; Wallden et al. 2016; Kiviet et al. 2014; Taheri-Araghi et al. 2015) and yeasts (Nobs & Maerkl 2014; Soifer, Robert & Amir 2016) mostly fell around the line of λ = 0.5 indicative of a near-adder behavior (note that, as expected, the yeast *S. Pombe* showed a better than adder behavior, but not a perfect sizer, because larger cells still have a minimal growth period (Fantes 1977)). These cells all showed a consistent degree of correction via time modulation (*θ* > 0). A small subset also showed positive growth rate modulation (*γ* > 0) Figure 5E, green box), but many also had instead a strong negative growth rate modulation (i.e. growth rate modulations contributed to noise in size instead of correcting it, Figure 5E blue box).

### The near-adder behavior emerges from a variety of couplings between growth and time modulations

Most mammalian cells displayed volume homeostasis close to an adder behavior (all points fell clustered around the line representing λ = 0.5, Figure 5F), consistent with the plot shown in Figure 2B. Data obtained from Son et al (Son et al. 2012) on L1210 showed that these cells were also adders, with mostly growth rate modulation (in accordance with the results of that study), but also some level of time modulation, possibly explained by the negative correlation between G1 duration and early growth speed observed in these cells (Son et al. 2012) (Figure 5F, yellow star). For both mammalian cells and bacteria, no dataset showed a negative time modulation (bigger cells at birth having a longer cell division cycle), meaning that time modulation always contribute to homeostasis, although to a lesser extent in mammalian cells than in yeasts and bacteria. Negative growth rate modulation (larger cells with a faster exponential growth rate than smaller cells at birth) was rarer in mammalian cells than in bacteria, but nevertheless observed in some cases (for Raji cells, Figure 5F, pink circle). This means that in mammalian cells, contrary to yeasts and bacteria, growth rate modulation always contributes to homeostasis and does it to a larger extent. Our analysis method, by providing a summarized overview of a large dataset comprising various cell types and culture conditions, demonstrated the generality of the phenomenological adder behavior, and also revealed the diversity of the underlying homeostatic mechanisms with different coupling of growth rate and timing modulation. Such diversity was observed even for a given cell line depending on the growth conditions (datasets from bacteria) or initial size (results from Roscovitine-induced large HeLa cells).

## Discussion

Due to technical limitations, our understanding of size homeostasis in mammalian cells derives in large part from indirect evidence (Conlon et al. 2001; Ian Conlon and Raff 2003; Echave, Conlon, and Lloyd 2007; Tzur et al. 2009; Sung et al. 2013; R. Kafri et al. 2013; Dolznig et al. 2004), nurturing controversies which have proven hard to resolve. To make progress in this area, we have developed a new method, FXm (Cadart et al. 2017; Zlotek-Zlotkiewicz et al. 2015), to follow the volume of single cultured mammalian cells over long periods of time, produced new datasets from direct measurements of freely-growing and dividing cells, and introduced conceptual tools to quantify the results.

In mammalian cells, a recent study provided direct evidence that size control relies on modulation of G1 duration (Varsano et al. 2017). Authors show that G1 duration negatively correlates with volume at birth, up to a point above which large cells cycle in a minimal time independent of initial size. Our experiment on control cells and artificially induced large cells nicely agree with this observation (Figure 3D-E and 4B). Remarkably, our description of size control in G1 combined with that of Varsano and colleagues is compatible with the most recent advances in the study of the cell cycle networks acting in G1 that identified two sub-periods in G1: a first period of variable length that involves pRb-E2F activation (Cappell et al. 2016; Barr et al. 2016), and a second period with a constant duration of 4 hours ending with the inactivation of APC-Cdh1 (Cappell et al. 2016). We envision that future research, combining tracking of key regulators of the cell cycle with volume measurement at the single-cell level (Schmoller et al. 2015) will help identify the long-hypothesized G1 size-checkpoints in mammalian cells (Fisher 2016).

Our dataset also provides direct evidence in support of the previously hypothesized role for growth rate modulations in size homeostasis (Kafri et al. 2013; Tzur et al. 2009). In particular, experiments on Roscovitine-induced abnormal large cells show that such cells grow on average exponentially (Figure 4D and S4D-E), do not adapt G1 duration to initial size (Figure 4B) and yet maintain an homeostatic behavior (Figure 4C). This is thanks to an adaptation of the exponential growth rate to the volume at birth (Figure 4E). We find that cells typically grow with a growth speed that increases as a function of time (and volume), but that the average growth rate (the slope of the growth speed as a function of volume) is smaller for cells born bigger, which is a new type of homeostatic behavior.

The pattern of growth of animal cells has been subject to debate, with contradicting conclusions as to whether they grow linearly (Conlon & Raff 2003; Varsano et al. 2017) or exponentially (Tzur et al. 2009; Son et al. 2012; Kafri et al. 2013; Mir et al. 2011; Sung et al. 2013). An illustration of how the growth mode has a strong impact on size control is the difference in effective homeostatic behavior we observed between our experiments and those from the recent work of Varsano and colleagues (Varsano et al. 2017) despite cells showing the same pattern of adaptation of G1 duration to size at birth. In the study by Varsano and coworkers, cells, which were confined in microchannels, grew linearly (an observation we reproduced with our microchannels experiments, see Figure S2A) and the combination of linear growth with G1 adaptation to volume at birth lead to a sizer-like behavior for small cells. In our study, unconfined cells in the FXm device grew nearly exponentially on average (HT29: Figure S1G, HeLa: Figure 4D) and thus, the same modulation of G1 duration lead to a weaker effective homeostatic behavior, whose strength was half way between an adder and a sizer (Figure 3B) for HT29 cells and close to an adder for HeLa cells (Figure 4C).

When considering growth trajectories, which is a unique feature of our dataset, we observed that individual cells could display complex growth behaviors, alternating plateaus and growth phases (Figures S4G-I), not clearly correlated with cell cycle stage events, even if plateaus were often seen in early G1. Growth in volume was on average faster than linear, when many cells were averaged (Figure S1G, 4D and S4D-E) but was, at the single cell level, highly modulated. A possible interpretation of this observation is that the average exponential (or faster than linear) growth is the result of multiple compensatory processes. The factors that could modulate growth rate at the single-cell level in a size-dependent manner are to date unknown and could involve, as recently hypothesized, the limitations of protein synthesis rate for large cell sizes (M. Kafri et al. 2016), nonlinear metabolic scaling with cell size (Miettinen & Bjorklund 2016) or physical constraints on volume growth via the addition of surface area (Glazier 2014), or dynamic changes in cell/substrate adhesion, cell spreading and cortical tension.

Our unbiased mathematical framework quantifies, for all cell types and all growth conditions, the respective contributions of growth and time modulation to the effective homeostatic behavior (Figure 5E-F). This analysis allowed us to compare the homeostatic behavior of widely different cells, and revealed a global similarity, but also striking differences between mammalian cells and unicellular organisms. First, the near-adder seems to be the most general behavior, found across kingdoms. In our data, four out of five different cell lines tested work as near adders, meaning that the added volume across a cell cycle was independent of the volume at birth. Such behavior has been observed in a variety of unicellular organisms, from bacteria (Campos et al. 2014; Taheri-Araghi et al. 2015; Soifer, Robert & Amir 2016) to budding yeast (Soifer, Robert & Amir 2016). However, the apparent universality of the adder at the phenomenological level may mask a more complex picture where several regulatory mechanisms acting in parallel or sequentially might be at play (Osella et al. 2017). Consequently, we find it unlikely that a single molecular ‘adder circuit’ would be working for mammalian cells. Second, and potentially more important, only mammalian cells combine both time and growth regulation to achieve homeostasis (in various ways depending on the cell type or the initial size for the same cell type, see Figure 5F). Unicellular organisms seem to only modulate timing. Environmentally-dictated changes in growth rate are widely regarded as a central parameter for cell size homeostasis in multicellular organisms (Roberts & Lloyd 2012; Lloyd 2013; Grewal & Edgar 2003). Thus, we surmise that the flexibility in the pattern of growth, characterizing mammalian cells only, may have to do with the acquisition of controlled and coordinated growth in tissues, which requires cells to respond quickly and efficiently to several joint environmental cues.

## Authors contributions

C.C. and S.M. conducted the experiments, S.M. optimized and designed the FXm chambers, E.T. and R.A. designed, produced and characterized the molds for the chambers, M.C-L. and J.G. developed the theoretical framework, B.B. helped conceive of project, helped supervise early part, helped with text, C.C. conducted the analysis, M.C-L. helped with data analysis, C.C. and M.C-L helped with the manuscript preparation, C.C. and M.P. designed the experiment, M.P. wrote the paper and supervised the work.

## Acknowledgements

We would like to thank the Gerlich lab for sharing the HeLa-MP cell line, Helen K. Matthews, Nunu Mchedlichvili and Ewa Zlotek-Zlotkiewicz, members of the Piel lab and the Perez lab for scientific and technical advices, Camille Blakeley and Charlotte Pirot for preliminary works as undergrad students, Thomas Wyatt for help on revising the manuscript, Isabel Brito for advices on the statistical analysis, the imaging platform from the Institut Curie PICT-IBiSA, the UMR 168 clean room facility and the IPGG plateform. We also greatly acknowledge Jan Skotheim for critical reading of the manuscript and in depth comments on our work. CC acknowledges support from the Fondation pour la Recherche Médicale (FDT20160435078) and the Ligue Nationale contre le Cancer for funding. MCL acknowledges support from the International Human Grantier Science Program Organization, grant RGY0070/2014. BB acknowledges Cancer Research UK programme grant for support: C1529/A17343. This work was supported by a LABEX IPGG grant to R.A., by an ERC consolidator grant (311205 PROMICO) to M.P., by an ANR grant to M.P. (ANR-14-CE11-0009-03, CellSize).

## References

Adiciptaningrum, A. et al., 2015. Stochasticity and homeostasis in the E. coli replication and division cycle. Scientific Reports, 5, p. 18261. Available at: http://www.nature.com/articles/srep18261 [Accessed January 24, 2017].

Amir, A., 2014. Cell Size Regulation in Bacteria. Physical Review Letters, 112(20), p.208102. Available at: http://link.aps.org/doi/10.1103/PhysRevLett.112.208102 [Accessed October 17, 2014].

Barr, A.R. et al., 2016. A Dynamical Framework for the All-or-None G1/S Transition. Cell Systems, 2(1), pp.27–37.

Cadart, C. et al., 2014. Exploring the function of cell shape and size during mitosis. Developmental Cell, 29(2).

Cadart, C. et al., 2017. Fluorescence eXclusion Measurement of volume in live cells. Methods in cell biology, 139(Cell polarity and morphogenesis), pp.103–120. Available at: http://www.elsevier.com/copyright.

Campos, M. et al., 2014. A Constant Size Extension Drives Bacterial Cell Size Homeostasis. Cell,159(6), pp.1433–1446. Available at: http://linkinghub.elsevier.com/retrieve/pii/S0092867414014998 [Accessed December 4, 2014].

Cappell, S.D. et al., 2016. Irreversible APCCdh1 Inactivation Underlies the Point of No Return for Cell-Cycle Entry. Cell, 166(1), pp.167–180.

Conlon, I. et al., 2001. Extracellular control of cell size. Nature cell biology, 3(October 2001), pp.3–7.Available at: http://discovery.ucl.ac.uk/186032/.

Conlon, I. & Raff, M., 2003. Differences in the way a mammalian cell and yeast cells coordinate cell growth and cell-cycle progression. Journal of biology, 2(1), p.7. Available at: /pmc/articles/PMC156598/?report=abstract [Accessed January 7, 2015].

Conlon, I. & Raff, M., 1999. Size control in animal development. Cell, 96(2), pp.235–244.

Deforet, M., Van Ditmarsch, D. & Xavier, J., 2015. Cell-Size Homeostasis and the Incremental Rule in a Bacterial Pathogen. Biophysical Journal, 109(3), pp.521–528.

Dolznig, H. et al., 2004. Evidence for a size-sensing mechanism in animal cells. Nature cell biology, 6(9), pp.899–905. Available at: http://www.ncbi.nlm.nih.gov/pubmed/15322555 [Accessed January 5, 2015].

Echave, P., Conlon, I.J. & Lloyd, A.C., 2007. Cell Size Regulation in Mammalian Cells. Cell Cycle, 6(2), pp.218–224. Available at: http://www.tandfonline.com/doi/abs/10.4161/cc.6.2.3744 [Accessed January 8, 2015].

Fantes, P.A., 1977. Control of cell size and cycle time in Schizosaccharomyces pombe. J Cell Sci, 24, pp.51–67. Available at: http://jcs.biologists.org/cgi/reprint/24/1/51.

Fisher, R.P., 2016. Getting to S: CDK functions and targets on the path to cell-cycle commitment. F1000Research, 5, p.2374. Available at: http://www.ncbi.nlm.nih.gov/pubmed/27746911.

Ginzberg, M.B., Kafri, R. & Kirschner, M., 2015. On being the right (cell) size. Science, 348(6236), pp.1245075–1245075. Available at: http://www.sciencemag.org/cgi/doi/10.1126/science.1245075.

Glazier, D., 2014. Metabolic Scaling in Complex Living Systems. Systems, 2(4), pp.451–540. Available at: http://www.mdpi.com/2079-8954/2/4/451/ [Accessed March 1, 2017].

Godin, M. et al., 2010. Using buoyant mass to measure the growth of single cells. Nature methods,7(5), pp.387–90. Available at: http://www.pubmedcentral.nih.gov/articlerender.fcgi?artid=2862099& tool=pmcentrez& render> type=abstract [Accessed December 3, 2014].

Grewal, S.S. & Edgar, B. a, 2003. Controlling cell division in yeast and animals: does size matter? Journal of biology, 2(1), p.5.

Grilli, J. et al., 2017. Relevant parameters in models of cell division control. Physical Review E - Statistical, Nonlinear, and Soft Matter Physics, 32411(95). Available at: http://arxiv.org/abs/1606.09284.

Grover, W.H. et al., 2011. Measuring single-cell density. Proceedings of the National Academy of Sciences of the United States of America, 108(27), pp.10992–6. Available at: http://www.pubmedcentral.nih.gov/articlerender.fcgifiartid=3131325&tool=pmcentrez&rendertype=abstract [Accessed December 5, 2014].

Harris, L.K. & Theriot, J.A., 2016. Relative Rates of Surface and Volume Synthesis Set Bacterial Cell Size. Cell, 165(6), pp.1479–1492. Available at: http://linkinghub.elsevier.com/retrieve/pii/S0092867416306481.

Hartwell, L. & Unger, M., 1977. Unequal Division in the Control Division and Its Implications for the Control of Cell Division. The Journal of cell biology, 75, pp.422–435.

Ho, P. & Amir, A., 2015. Simultaneous Regulation of Cell Size and Chromosome Replication in Bacteria. Frontiers in Microbiology, 6(July), pp.1–10.

Horvath, A. et al., 2016. Cell length growth patterns in fission yeast reveal a novel size control mechanism operating in late G2 phase. Biology of the Cell, 108(9), pp.259–277.

Iyer-Biswas, S. et al., 2014. Scaling laws governing stochastic growth and division of single bacterial cells. Proceedings of the National Academy of Sciences, p.1403232111-. Available at: http://www.pnas.org/content/early/2014/10/23/1403232111.abstract.html?etoc.

Johnston, G.C., Singer, R. a & McFarlane, S., 1977. Growth and cell division during nitrogen starvation of the yeast Saccharomyces cerevisiae. Journal of bacteriology, 132(2), pp.723–30. Available at: http://www.pubmedcentral.nih.gov/articlerender.fcgi?artid=221916&tool=pmcentrez&rendertype=abstract.

Jorgensen, P. & Tyers, M., 2004. How cells coordinate growth and division. Current biology?: CB,14(23), pp.R1014–27. Available at: http://www.ncbi.nlm.nih.gov/pubmed/15589139 [Accessed September 1, 2014].

Jun, S. & Taheri-Araghi, S., 2015. Cell-size maintenance: Universal strategy revealed. Trends in Microbiology, 23(1), pp.4–6. Available at: http://dx.doi.org/10.1016/j.tim.2014.12.001.

Kafri, M. et al., 2016. Rethinking cell growth models. FEMS Yeast Research, 16(7), pp.1–13.

Kafri, R. et al., 2013. Dynamics extracted from fixed cells reveal feedback linking cell growth to cell cycle. Nature, 494(7438), pp.480–483. Available at: http://dx.doi.org/10.1038/nature11897.

Kennard, A.S. et al., 2016. Individuality and universality in the growth-division laws of single E. Coli cells. Physical Review E - Statistical, Nonlinear, and Soft Matter Physics, 93(1), pp.1–18.

Killander, D. & Zetterberg, A., 1965. quantitative cytochemical studies on interphase growth II derivation of synthesis curves from the distribution of DNA, RNA and mass values of individual mouse fibroblasts in vitro. Experimental Cell Research, 39, pp.22–32.

Kiviet, D.J. et al., 2014. Stochasticity of metabolism and growth at the single-cell level. Nature,514(7522), pp.376–379. Available at: http://dx.doi.org/10.1038/nature13582.

Lancaster, O.M. et al., 2013. Mitotic rounding alters cell geometry to ensure efficient bipolar spindle formation. Developmental cell, 25(3), pp.270–83. Available at: http://www.ncbi.nlm.nih.gov/pubmed/23623611 [Accessed December 19, 2014].

Lloyd, A.C., 2013. The regulation of cell size. Cell, 154(6), pp.1194–205. Available at: http://www.ncbi.nlm.nih.gov/pubmed/24034244 [Accessed July 18, 2014].

Meijer, L. & Raymond, E., 2003. Roscovitine and Other Purines as Kinase Inhibitors. From Starfish Oocytes to Clinical Trials Roscovitine and Other Purines as Kinase Inhibitors. From Starfish Oocytes to Clinical Trials. Acc. Chem. Res, 36(6), pp.417–425.

Miettinen, T.P. & Bjorklund, M., 2016. Cellular Allometry of Mitochondrial Functionality Establishes the Optimal Cell Size. Developmental Cell, 39(3), pp.370–382.

Mir, M. et al., 2011. Optical measurement of cycle-dependent cell growth. Proceedings of the National Academy of Sciences of the United States of America, 108(32), pp.13124–9. Available at: http://www.pubmedcentral.nih.gov/articlerender.fcgifiartid=3156192&tool=pmcentrez&render type=abstract [Accessed November 18, 2014].

Mitchison, J.M., 2003. Growth During the Cell Cycle. International Review of Cytology, 226, pp.165–258. Available at: http://www.sciencedirect.com/science/article/pii/S0074769603010040.

Nobs, J.-B. & Maerkl, S.J., 2014. Long-term single cell analysis of S. pombe on a microfluidic microchemostat array. M. Polymenis, ed. PloS one, 9(4), p.e93466. Available at: http://dx.plos.org/10.1371/journal.pone.0093466 [Accessed February 8, 2017].

Osella, M., Nugent, E. & Cosentino Lagomarsino, M., 2014. Concerted control of Escherichia coli cell division. Proceedings of the National Academy of Sciences of the United States of America, 111(9), pp.3431–5. Available at: http://www.pubmedcentral.nih.gov/articlerender.fcgifiartid=3948223&tool=pmcentrez&rendertype=abstract [Accessed December 22, 2014].

Osella, M., Tans, S.J. & Cosentino Lagomarsino, M., 2017. Step by Step, Cell by Cell: Quantification of the Bacterial Cell Cycle. Trends in Microbiology, xx, pp.1–7. Available at: http://linkinghub.elsevier.com/retrieve/pii/S0966842X16302001.

Pan, K.Z. et al., 2014. Cortical regulation of cell size by a sizer cdr2p. eLife, 2014(3), p.e02040. Available at: http://www.ncbi.nlm.nih.gov/pubmed/24642412 [Accessed January 30, 2017].

Park, K. et al., 2010. Measurement of adherent cell mass and growth. Proceedings of the National

Academy of Sciences of the United States of America, 107(48), pp.20691–6. Available at: http://www.pubmedcentral.nih.gov/articlerender.fcgifiartid=2996435&tool=pmcentrez&rendertype=abstract [Accessed January 5, 2015].

Popescu, G. et al., 2014. New technologies for measuring single cell mass. Lab on a chip, 14(4), pp.646–52. Available at: http://www.ncbi.nlm.nih.gov/pubmed/24322181.

Roberts, S. a. & Lloyd, A.C., 2012. Aspects of cell growth control illustrated by the Schwann cell. *Current Opinion in Cell Biology*, 24(6), pp.852–857. Available at: http://dx.doi.org/10.1016/j.ceb.2012.10.003.

Sakaue-Sawano, A. et al., 2008. Visualizing Spatiotemporal Dynamics of Multicellular Cell-Cycle Progression. Cell, 132(3), pp.487–498.

Sauls, J.T., Li, D. & Jun, S., 2016. Adder and a coarse-grained approach to cell size homeostasis in bacteria. Current Opinion in Cell Biology, 38, pp.38–44.

Schmoller, K. et al., 2015. Dilution of the cell cycle inhibitor Whi5 controls budding yeast cell size. Nature, (in press).

Soifer, I., Robert, L., Amir, A., et al., 2016. Single-Cell Analysis of Growth in Budding Yeast and Bacteria Reveals a Common Size Regulation Report Single-Cell Analysis of Growth in Budding Yeast and Bacteria Reveals a Common Size Regulation Strategy. Current Biology, 26(3), pp.356– 361. Available at: http://dx.doi.org/10.1016/j.cub.2015.11.067.

Soifer, I., Robert, L. & Amir, A., 2016. Single-cell analysis of growth in budding yeast and bacteria reveals a common size regulation strategy. Current Biology, 26(3), pp.356–361.

Sompayrac & Maaloe, O., 1973. Autorepressor Model for Control of DNA Replication. Nature: new biology, 241.

Son, S. et al., 2012. Direct observation of mammalian cell growth and size regulation. Nature Methods, 9(9), pp.910–912.

Son, S. et al., 2015. Resonant microchannel volume and mass measurements show that suspended cells swell during mitosis. Journal of Cell Biology, 211(4), pp.757–763.

Sung, Y. et al., 2013. Size homeostasis in adherent cells studied by synthetic phase microscopy.*Proceedings of the National Academy of Sciences of the United States of America*, 110(41), pp.16687–92. Available at: http://www.pubmedcentral.nih.gov/articlerender.fcgi?artid=3799364&tool=pmcentrez&rendertype=abstract [Accessed January 5, 2015].

Sveiczer, A., Novak, B. & Mitchison, J.M., 1996. The size control of fission yeast revisited. Journal of cell science, 109, pp.2947–2957.

Taheri-Araghi, S. et al., 2015. Cell-Size Control and Homeostasis in Bacteria. Current Biology, 25(3), pp.385–391. Available at: http://linkinghub.elsevier.com/retrieve/pii/S0960982214015735.

Di Talia, S. et al., 2007. The effects of molecular noise and size control on variability in the budding yeast cell cycle. Nature, 448(7156), pp.947–51. Available at: http://www.ncbi.nlm.nih.gov/pubmed/17713537 [Accessed September 1, 2014].

Turner, J.J., Ewald, J.C. & Skotheim, J.M., 2012. Cell size control in yeast. Current Biology, 22(9).

Tzur, A. et al., 2009. Cell growth and size homeostasis in proliferating animal cells. Science (New York, N.Y.), 325(5937), pp.167–71. Available at: http://www.pubmedcentral.nih.gov/articlerender.fcgifiartid=2905160&tool=pmcentrez&rendertype=abstract [Accessed January 5, 2015].

Varsano, G. et al., 2017. Probing Mammalian Cell Size Homeostasis by Article Probing Mammalian Cell Size Homeostasis by Channel-Assisted Cell Reshaping. CellReports, 20(2), pp.397–410. Available at: http://dx.doi.org/10.1016/j.celrep.2017.06.057.

Voorn, W.J. & Koppes, L.J.H., 1998. Skew or third moment of bacterial generation times. Archives of Microbiology, 169(1), pp.43–51.

Wallden, M. et al., 2016. The synchronization of replication and division cycles in individual E. coli cells (in press). Cell, pp.729–739.

Wang, P. et al., 2010. Robust growth of escherichia coli. Current Biology, 20(12), pp.1099–1103.Available at: http://dx.doi.org/10.1016/j.cub.2010.04.045.

Willis, L. et al., 2016. Cell size and growth regulation in the Arabidopsis thaliana apical stem cell niche. Proceedings of the National Academy of Sciences of the United States of America, p.201616768. Available at: http://www.ncbi.nlm.nih.gov/pubmed/27930326.

Wood, E. & Nurse, P., 2013. Pom1 and cell size homeostasis in fission yeast. Cell cycle (Georgetown, Tex.), 12(19), pp.3228–36. Available at:http://www.pubmedcentral.nih.gov/articlerender.fcgi?artid=3865018&tool=pmcentrez&rendertype=abstract [Accessed January 7, 2015].

Yu, F.B. et al., 2017. Long-term microfluidic tracking of coccoid cyanobacterial cells reveals robust control of division timing. BMC Biology, 15(1), p.11. Available at: http://bmcbiol.biomedcentral.com/articles/10.1186/s12915-016-0344-4.

Zielke, N. et al., 2011. Control of Drosophila endocycles by E2F and CRL4CDT2. Nature, 480(7375), pp.123–127.

Zlotek-Zlotkiewicz, E. et al., 2015. Optical volume and mass measurements show that mammalian cells swell during mitosis. The Journal of Cell Biology, 211(4), pp.765–774. Available at: http://www.jcb.org/cgi/doi/10.1083/jcb.201505056.

